# A complete adult *Drosophila melanogaster* brain tracheolar network

**DOI:** 10.64898/2026.11.09.765327

**Authors:** Haneul Cho, Thea Ng, Kira Schoenberg, Sommer Huntress, Jimin Suh, Athena Manandhar, Shuojia Lin, Bethelehem M. Woldetsadik, Rachel S. Kang, Mai Bui, Louisa M. Zebrowski, Fiona Reynolds-Cornell, Elizabeth Agbedun, Yaslyn Cruz, Caroline A. Lewis, Kesshni Bhasiin, Keying Lin, Carmela Mooney, Andrew M. Dacks, Philipp Schlegel, Davi D. Bock, Kenneth J. Colodner

## Abstract

The tracheal system of the fruit fly, *Drosophila melanogaster*, is a complex network of epithelial tubules that supplies oxygen and removes carbon dioxide throughout the organism, with gases entering and exiting through spiracles along the body. While tracheal development and patterning in the embryonic and larval stages have been extensively studied, particularly in the body, much less is known about tracheal and tracheolar organization within the adult brain. To address this gap, we manually traced every tracheole in the brain of a female adult *Drosophila* using the Full Adult Fly Brain (FAFB) open-access electron microscopy dataset. We reconstructed 117 tracheoles that branch within the parenchyma, terminating in 1,376 endings without forming an interconnected network. Joint analysis of these skeleton reconstructions with volumetric mesh segmentations from the FlyWire connectome revealed that tracheole volume fraction is heterogeneous across neuropils and largely symmetric in overall tracheation between left and right hemispheres. We observe that synapses are not preferentially positioned near tracheoles, and are instead slightly depleted from a region approximately 15 µm around tracheoles, relative to other points along the same neurons. We also quantify the relative proximity of neurotransmitter-specific synapses and cables to tracheoles across neuropils. This detailed anatomical description of the tracheolar network establishes a reference for future studies examining how neuronal patterning, activity, and metabolic demand couple with tracheole development and function.

## Introduction

The tracheal system of *Drosophila melanogaster* is a branching network of gas-filled epithelial tubes that enables cellular respiration by facilitating oxygen delivery and carbon dioxide removal throughout the organism (Wigglesworth, 1942). This network originates at spiracles in the lateral thorax and abdomen, and supports the metabolic demands of all tissues at every developmental stage. During embryogenesis, the tracheal system is established with bilateral symmetry, where it displays a stereotyped, interconnected pattern across the metameric units of the body. The embryonic tracheal network is subsequently remodeled during larval development through enlargement and elongation of tracheal tubes, and again during metamorphosis, when adult trachea precursor cells (tracheoblasts) reconstruct the larval network to form adult-specific tracheal branches (Hayashi & Kondo, 2018). The intrinsic and extrinsic molecular mechanisms that control *Drosophila* body tracheal development have been well-characterized, offering a rich model for studying respiration and tubulogenesis (Bossen et al., 2023; Ehrhardt et al., 2022).

In addition to supplying the body’s respiratory needs, the tracheal network also supports gas exchange in the *Drosophila* nervous system. During embryogenesis, the ventral nerve cord becomes tracheated via ganglionic branches arising from the dorsal trunk tracheae (Englund et al., 1999), while the embryonic brain receives tracheae through the cervical (cerebral) branch tracheal network that originates from the first segmental trachea in the second thoracic segment (Hartenstein et al., 2008; Pereanu et al., 2007). These primary cervical branches differentiate into dorsal anterior and posterior trunks, which in turn give rise to lateral, ventral, and medial trunks that infiltrate and tracheate the larval and pupal brain (Hartenstein et al., 2008; Whitten, 1957). In the adult, tracheoblasts again remodel the dorsal cervical tracheae, which dilate into air sacs that encircle the brain. From these sacs, numerous finer branches of trachea, termed tracheoles, extend into both hemispheres and supply the adult brain with oxygen (Bate & Martinez Arias, 1993).

While the general anatomy of external trachea and air sacs in the adult brain is fairly stereotyped, the finer, internal tracheolar branching patterns are more variable and less well understood (Hartenstein et al., 2008). A detailed analysis of the adult brain tracheolar network is hampered by the fact that the finest tracheoles responsible for directly tracheating the brain are < 1 µm in diameter, and thus difficult to resolve using conventional fluorescence microscopy. Moreover, the expression of common tracheal genetic reporters (e.g. *btl*) is markedly reduced in adult stages, limiting the utility of genetic tools that have been instrumental in developmental tracheal mapping (Bossen et al., 2023). As a result, several basic questions about the organization of the adult brain tracheolar network remain to be determined. It is unclear whether tracheation is uniformly distributed throughout the brain, and whether tracheolar patterning is symmetric across brain hemispheres. Moreover, tracheoles typically infiltrate and come to terminal endings within tissue (Wigglesworth, 1942), but whether this anatomy holds for the fine tracheoles that penetrate the adult brain, as opposed to an interconnected network akin to anastomosing vertebrate vasculature, has not been determined.

To advance our understanding of the structure of the adult *Drosophila* brain tracheolar network, we manually traced every tracheole in a high-resolution, open-access electron microscopy dataset of an adult female *Drosophila* brain (Zheng et al., 2018). We characterize the full neuroanatomical distribution, branching architecture, and morphological features of this network, and integrate our manual tracing ground truth with volumetric segmentation data (FlyWire project; Dorkenwald et al., 2024; Schlegel et al., 2024) to assess tracheole volume fraction and proximity to synapses and neuronal processes throughout the brain (Buhmann et al., 2021; Eckstein et al., 2024; Yu et al., 2025). This comprehensive analysis establishes a foundational map of the adult *Drosophila* brain tracheolar system and provides a resource for future interrogations into fine-scale tracheole stereotypy, in addition to the structural and functional interplay between tracheolar networks and brain activity.

## Materials and Methods

### Tracheole Tracing

Tracheole skeletons were manually traced in the Full Adult Fly Brain (FAFB) dataset of a 7-day-old female fly of the genotype: [iso] w^1118^ x [iso] Canton-S G1 (Zheng et al., 2018). Tracing was performed using the Collaborative Annotation Toolkit for Massive Amounts of Image Data software (CATMAID; RRID: SCR_006278) (Saalfeld et al., 2009). Trained tracers placed nodes at the center of tracheole lumens, and each skeleton was reviewed for completeness and accuracy by at least one of three expert reviewers.

The root of each tracheole skeleton was placed at the point of entry (infiltration point) into the brain, which was defined as the center of the lumen at the neural lamella/glial sheath barrier. For one tracheole lacking a visible infiltration point, the root was placed at the widest segment nearest the brain’s exterior. Tracheoles external to the brain, and those within the optic laminae, were excluded from analysis.

To verify completeness of the tracing effort, two independent tracers reviewed electron micrographs at every 250th z-section across the entire brain volume. All tracheole lumens in these sections were examined and the presence of placed nodes at lumen centers was confirmed.

### Proofreading and segmentation in FlyWire

Manually traced tracheole skeletons were imported into FlyWire (RRID:SCR_019205) and used to identify corresponding segmented meshes. Where necessary, disparate supervoxels representing the same tracheole were merged, and contaminating adjacent glial cells and neuronal elements were manually split from the tracheolar mesh. Tracheole supervoxels that could not be merged without including non-tracheole structures were excluded from the tracheole mesh. The proofread tracheole meshes and CATMAID tracheole skeletons were analyzed using the python libraries: Neuron Analysis and Visualizations (NAVis) (Bates et al., 2020) (https://navis-org.github.io/navis) and pymaid: a Python-CATMAID Interface (https://github.com/navis-org/pymaid). A link to the proofread meshes can be found here, after authorization with a Google account through FlyWire: https://ngl.flywire.ai/?json_url=https://globalv1.flywire-daf.com/nglstate/6146478859354112.

### Tracheole volume fraction quantification

Tracheole volume fraction per neuropil volume, was calculated by dividing the volume of all tracheole meshes contained within a neuropil mesh volume, by the volume of the neuropil, and multiplied by 100. This volume fraction was quantified from FlyWire segmentation voxels and FlyWire brain-region meshes, and each CATMAID tracheole skeleton was linked to a corresponding FlyWire mesh.

The retained segmentation was represented by voxel centers at a resolution of 0.064 × 0.064 × 0.040 µm. To assign these tracheole voxels to individual brain regions, voxels outside each region’s rectangular bounding box were removed as a computational prefilter. The remaining candidate voxel centers were then tested against the actual closed FlyWire brain-region mesh using a point-in-mesh test. Voxels inside the mesh were counted toward that region. Regional tracheole volume was calculated as the number of retained voxels multiplied by the volume of one voxel, and tracheole volume fraction was calculated as 100 × Vtracheole / Vregion. For super-region analyses, tracheole volumes and brain-region volumes were summed separately across all constituent subregions before calculating the percentage. Super-regions were computed based on established neuroanatomical categories (Ito et al., 2014). Seven tracheoles located in the brain periphery were excluded from this volumetric analysis, as accurate FlyWire meshes corresponding to these tracheoles could not be obtained, due to an absence of segmentation at the outer periphery of the brain. Also, note that due to an inversion in FAFB acquisition (Schlegel et al., 2024), the anatomical left brain hemisphere is depicted on the viewer’s left.

### Infiltration point mapping

The node at the center of mass of the lumen at the neural lamella/glial sheath barrier was set as the skeleton root in CATMAID and tagged “Infiltration point” (n=116; one tracheole had no identifiable infiltration point and was excluded from this analysis). Because infiltration points lie at the brain surface, outside the boundaries of the annotated neuropil subvolumes, they could not be assigned to a brain region automatically. Each infiltration point was therefore assigned manually to a brain region by using the CATMAID 3D viewer, in which the tracheole skeleton was displayed together with the annotated neuropil volume meshes. The trajectory of each tracheole was followed from its root, and the first volume mesh it entered was recorded as the region of infiltration. Assignments were made by a single annotator, with discrepancies resolved by a second reviewer.

### Inter-tracheole spacing

To quantify spacing between neighboring tracheoles, the three-dimensional Euclidean distance from each CATMAID tracheole node to the nearest node belonging to a different tracheole was calculated. Nodes from the same tracheole were excluded as candidate neighbors. Distances were pooled across all tracheole nodes to characterize the distribution of inter-tracheole spacing.

### Strahler Analysis

Branching complexity was quantified from the parent-child topology of each CATMAID tracheole skeleton using Strahler number. Terminal nodes were assigned an order of 1. Moving from the terminal branches toward the root, a parent node was assigned the maximum order of its children when only one child had that order; when two or more children shared the maximum order, the parent order was increased by one. The Strahler number of each tracheole was defined as the order of its root node. Tracheoles were then grouped according to their resulting Strahler number.

### Tracheolar ends analysis

Terminal-end organization was analyzed across the 117 tracheole skeletons in CATMAID. Terminal ends were defined as distal CATMAID nodes with no children, excluding the CATMAID root node. Two nearest-neighbor analyses were performed. First, to quantify spacing between distinct tracheoles, each terminal end was paired with the nearest terminal end belonging to a different tracheole, with all ends from the same tracheole excluded. To quantify spacing within individual tracheoles, each terminal end was also paired with the nearest other terminal end on the same tracheole. Tracheoles with fewer than two terminal ends were excluded from this within-tracheole analysis. Distance distributions were summarized across individual terminal ends and, where indicated, by the mean nearest-end distance for each tracheole. Tracheoles were additionally classified by Strahler number to relate terminal-end organization to branching complexity.

### Tracheole Link Connector (TLC) count per volume

Membranous tethers connecting tracheole lumens on either side of the tether were identified and a node at the center point of the tether was tagged in CATMAID with ‘TLC_connection’. For each pruned skeleton within a defined subvolume, TLC_connection tags were identified using the CATMAID find_nodes function. The 3D coordinates (x, y, z) of each TLC_connection node were extracted using the get_node_location function and stored for spatial mapping. All TLC_connection nodes were visualized in the brain volume using the NAVis plot3d function.

### Synapse-to-tracheole proximity

Presynaptic contacts were obtained from the FAFB v783 Princeton synapse dataset (Yu et al., 2025), which improves upon the synapse predictions originally released with FlyWire (Buhmann et al., 2021). To represent the tracheolar network, CATMAID tracheole skeleton edges were resampled at 0.1-µm intervals to generate a dense three-dimensional centerline point cloud. For each presynaptic contact, the Euclidean distance from its presynaptic coordinate to the nearest sampled tracheole centerline point was calculated. Contacts in the left and right laminae were excluded from regional analyses. Contacts were assigned to anatomical neuropils and aggregated into super-regions as described above. For neurotransmitter-stratified analyses, each presynaptic neuron was assigned the neurotransmitter class with the highest prediction score, and contacts were retained when that score was ≥0.5.

### Synapse proximity to tracheole terminal ends

To separately assess proximity to distal tracheole endings, the 1,376 terminal-end coordinates identified from the 117 CATMAID tracheole skeletons were used as the spatial reference. For each presynaptic contact, the Euclidean distance to the nearest terminal end was calculated. Distance distributions were summarized across all contacts and separately by neurotransmitter class using the same neurotransmitter assignments described above. Neurotransmitter-specific terminal-end distance distributions were compared using a chi-square test on 1-µm distance bins, with distances ≥50 µm pooled into a tail bin. Cramér’s V was reported as the effect size.

### Anatomy-matched neurotransmitter proximity analysis

To determine whether neurotransmitter-specific synapse-to-tracheole distances differed from those expected from anatomical location alone, each neurotransmitter population was compared with an anatomy-matched null distribution. Within each super-region, the observed number of contacts for a given neurotransmitter was preserved, and the same number of contacts was sampled without replacement from all presynaptic contacts in that region, irrespective of neurotransmitter identity. This procedure generated the expected median distance for that neurotransmitter within each anatomical context. For the whole-brain analysis, matched samples were generated separately within each super-region and then combined before calculating the whole-brain median, preserving the observed regional occupancy of each neurotransmitter class. Observed-minus-expected median distances were used as the effect measure, with negative values indicating closer-than-expected and positive values indicating farther-than-expected synapse positions.

### Synaptic proximity to tracheole surfaces relative to local neuronal cable

Synaptic proximity to tracheole surfaces was tested by comparing presynaptic sites with matched non-presynaptic cable, or processes, from the same neuron. Neurons were assigned to the neurotransmitter class with the highest annotation score, and assignments with a score below 0.5 were excluded. Neurons were required to contain at least 20 synaptic contacts and at least 20 unique presynaptic sites. Because the numbers of eligible neurons differed substantially among neurotransmitter classes, up to 250 neurons were randomly selected from each class to reduce sampling imbalance; when fewer than 250 eligible neurons were available, all were retained. All unique presynaptic sites from the selected neurons were included in the subsequent analysis. The left and right laminae were excluded from the analysis.

Neuronal skeletons were resampled at 0.25-µm spacing to represent local neuronal cable. Presynaptic sites were required to lie within 5 µm of their corresponding neuronal skeleton. For each presynaptic site, three control positions were selected from non-presynaptic cable on the same neuron and 3-25 µm from the presynaptic site. When fewer than three eligible controls were available within this range, the search radius was extended to 40 µm. Cable positions within 1 µm of any known presynaptic site were excluded. Controls were sampled without replacement and were not reused for another presynaptic site on the same neuron.

For each presynaptic site and matched cable control, proximity to the tracheole system was measured as the shortest three-dimensional distance to the surface of the trimmed FlyWire tracheole meshes. For each matched set, synaptic displacement was calculated as the presynaptic distance minus the median distance of its three matched cable controls. Negative values indicate that the presynaptic site lies closer to the tracheole surface than the surrounding local cable, whereas positive values indicate that it lies farther away from it.

Presynaptic and matched-cable distance distributions were compared using empirical cumulative distribution functions (ECDFs). Negative ΔECDF values indicate a relative shift of presynaptic sites toward larger tracheole-surface distances, whereas positive values indicate a shift toward smaller distances. The signed Kolmogorov–Smirnov statistic was defined as the value of ΔECDF at the distance at which the absolute separation between the two distributions was maximal. The same analysis was performed separately for each neurotransmitter class.

To test whether synaptic displacement depended on the local tracheole proximity of the neuron itself, the median tracheole-surface distance of the three matched cable controls was used as the local cable-distance measure for each presynaptic site. Its relationship with synaptic displacement was quantified using Spearman rank correlation.

To test whether presynaptic sites were positioned closer to tracheoles than their matched controls, we used conditional logistic regression, which compares each presynaptic site with its own three matched cable controls. Distance to the tracheole surface was modeled as a continuous variable and scaled per 5 µm. Effects were expressed as the odds ratio per 5 µm closer to the tracheole surface, such that values greater than 1 indicate relative enrichment of presynaptic sites closer to trachea and values below 1 indicate relative displacement away from the trachea. The same matched analysis was performed for neurotransmitter and anatomical subgroups. Subgroup models were fitted only when at least 100 matched sets were available, and multiple comparisons were corrected using the Benjamini-Hochberg false-discovery-rate procedure.

To account for the contribution of multiple presynaptic sites from the same neuron, synaptic displacement was additionally summarized at the neuron level by taking the median displacement across all matched presynaptic sites from each neuron. The population effect was calculated as the median of these neuron-level values. Confidence intervals were obtained from 5,000 bootstrap resamples of neurons, and deviation from zero was tested using a Wilcoxon signed-rank test.

### Neurotransmitter-specific presynaptic arbor proximity

To determine whether neuronal cable from different neurotransmitter classes occupied different spatial relationships to the tracheolar network, presynaptic arbors were reconstructed from FlyWire neuronal skeletons for the same neurotransmitter-balanced neuron cohort used in the matched-cable analysis. Skeleton edges were resampled at 0.25-µm spacing, and analyzed presynaptic sites were mapped to the corresponding neuronal skeleton. For each neuron, the minimal connected skeleton subtree linking its analyzed presynaptic sites was retained as a presynaptic-arbor proxy. Neurons were retained only when at least 50% of analyzed presynaptic sites could be mapped within 5 µm of the skeleton. The distance from each cable position to the nearest surface of the trimmed FlyWire tracheole meshes was then calculated. Cable-distance distributions were summarized both by cable length and at the individual-neuron level, with one neuron used as the statistical unit for inferential comparisons among neurotransmitter classes. The same analysis was additionally performed using the entire neuronal skeleton as a whole-neuron sensitivity analysis.

### Bilateral tracheole symmetry

Bilateral tracheole symmetry was quantified using the 117 CATMAID tracheole skeletons transformed into FlyWire coordinates and the corresponding bilateral FlyWire neuropil meshes. Thirty-four left–right neuropil pairs were available for length and tracheation analyses; 33 yielded valid spatial-registration measurements. The left and right laminae were excluded from the bilateral region set. For each neuropil pair, tracheole length was measured separately on the left and right, and tracheole length asymmetry was calculated as (R−L)/(R+L), such that negative values indicate greater tracheole length on the left and positive values indicate greater length on the right.

To compare spatial organization after accounting for differences in the position and orientation of paired neuropils, the left neuropil and its associated tracheoles were first reflected across a common brain midline. A rigid registration was then estimated from the neuropil surfaces independently of the tracheole distributions. For each left–right neuropil pair, 15,000 points were sampled from each mesh surface. The reflected left surface was initially translated to align its centroid with that of the right surface and was then aligned by iterative closest-point registration. At each iteration, each transformed left-surface point was paired with its nearest point on the right surface. The closest 85% of correspondences were retained to reduce the influence of poorly matched surface regions, and the optimal rigid rotation and translation were estimated by the Kabsch algorithm. This procedure was repeated for up to 40 iterations and stopped when the change in median nearest-neighbor distance was less than 5 nm. The resulting rigid transformation was then applied to the reflected left-side tracheole centerline, placing it in the coordinate frame of the corresponding right neuropil.

Normalized spatial mismatch was calculated from the registered tracheole distributions. For each left-side centerline sample, the distance to its nearest right-side centerline sample was measured, and the calculation was repeated from right to left. The median nearest-neighbor distance was calculated in each direction, and the two medians were averaged to give a symmetric spatial mismatch. This value was normalized by the mean three-dimensional bounding-box diagonal of the paired left and right neuropil meshes to account for differences in region size. Local correspondence was quantified as the fraction of tracheole centerline samples whose nearest contralateral counterpart lay within 5 µm after registration. Correspondence was calculated separately from left to right and from right to left and averaged across the two directions.

### Visualization

3D visualizations were generated using NAVis.plot3d, plotly (https://plotly.com; RRID:SCR_013991), or by taking screenshots of CATMAID or FlyWire.

### Statistical Analyses

Statistical analyses were performed in Python with two-sided tests. Distributions were summarized by medians and interquartile ranges, and 95% bootstrap confidence intervals were estimated from 5,000 resamples. Group differences were assessed using Kruskal-Wallis tests with epsilon-squared effect sizes, followed where appropriate by Mann-Whitney U tests with rank-biserial effect sizes. Paired comparisons used Wilcoxon signed-rank tests with paired rank-biserial effect sizes, and associations were assessed using Spearman rank correlations.

Multiple comparisons were corrected using the Benjamini-Hochberg false-discovery-rate procedure. Tracheole volume scaling was assessed by linear regression in log-log space, including a test of the fitted slope against 1. For neurotransmitter-specific null analyses, observed distances were compared with 5,000 equal-sized samples drawn without replacement from the appropriate anatomical population; for the whole-brain analysis, these samples also preserved each neurotransmitter’s contact count within every super-region. Synapse-versus-cable comparisons used conditional logistic regression, with each presynaptic site and its matched controls treated as one set, with complementary neuron-level analyses based on median synaptic displacement. For analysis of neuronal cable, one neuron was treated as the statistical unit.

## Results

### Reconstruction of the adult brain tracheolar network

To determine how the brain’s gas exchange system is organized relative to its neuroanatomical compartments, we mapped the complete tracheolar network of an adult *Drosophila* brain in the Full Adult Fly Brain (FAFB) electron microscopy volume (Zheng et al., 2018). Because the neurons and synapses of this same brain have been comprehensively reconstructed (Dorkenwald et al., 2024; Schlegel et al., 2024), this approach allowed us to directly relate the position of every tracheole to the neuropil compartments, neurons, and synapses that comprise this brain. Using the web-based annotation software, CATMAID (Saalfeld et al., 2009), we manually skeletonized the tracheoles in the electron micrographs and identified 117 unique tracheoles spread throughout the brain (Figure 1A-C; Supplemental Video 1), noting regions of sparse and dense tracheation (Figure 1D, E). Total tracheolar cable length is 12.98 cm; in comparison, total neuronal path length within this same brain is 149.2 m (Dorkenwald et al., 2024). We imported the tracheole skeletons into the FlyWire platform (Dorkenwald et al., 2024; Schlegel et al., 2024) and retrieved intersecting segmented tracheolar meshes, which were manually proofread to remove incorrectly merged neurons and adjacent glial processes (Figure 1F, G). Glial processes were frequently found directly apposed to tracheoles and were often merged with them in the automated segmentation (Figure 1F). Manual proofreading resulted in a high-fidelity volumetric tracheolar network (Figure 1H).

**Figure 1.**
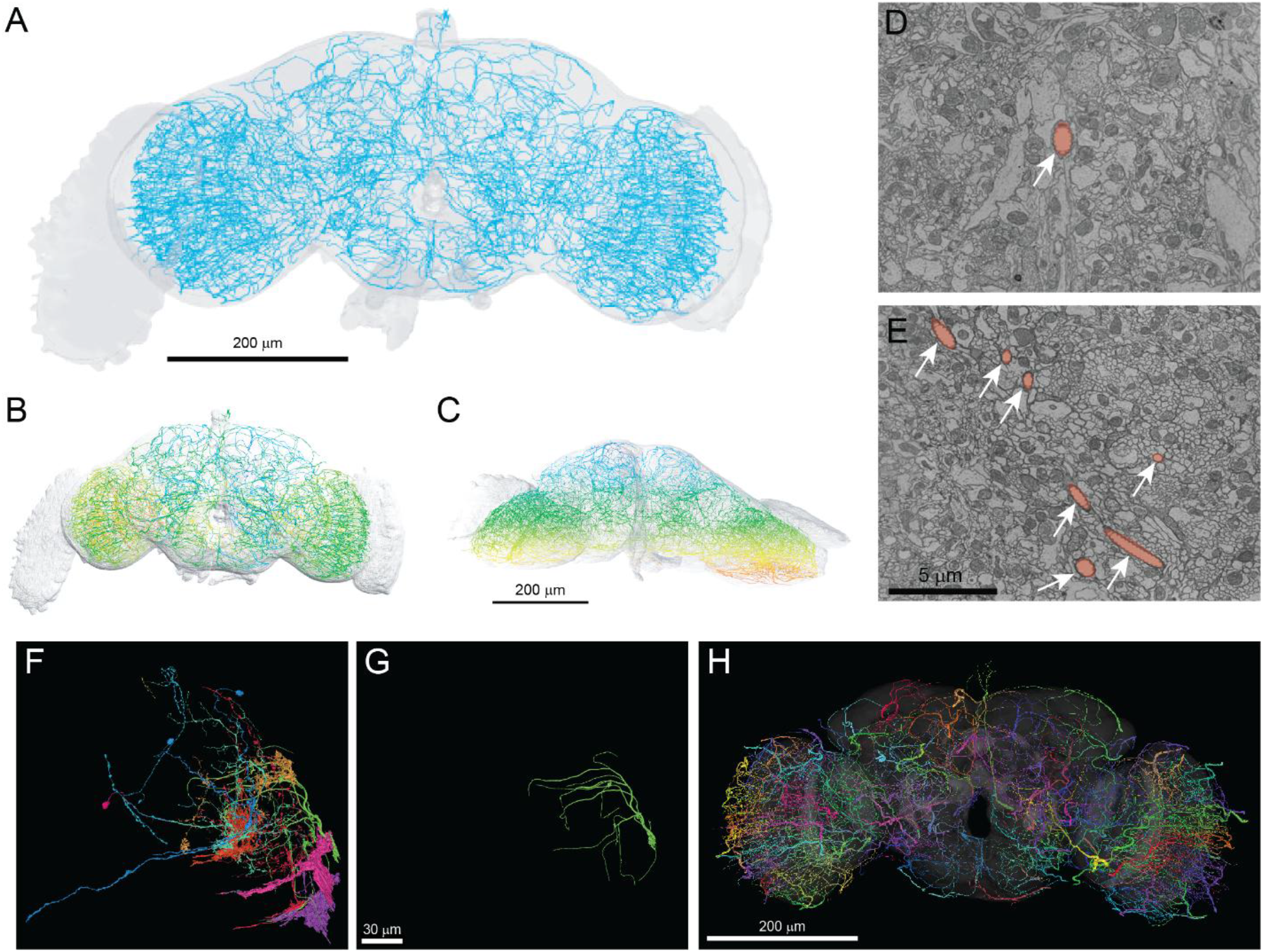
Neuroanatomical architecture of the adult *Drosophila* brain tracheolar network. (A) Frontal 3D view of the complete skeletonized brain tracheolar network. (B-C) Depth-mapping of all tracheoles, rainbow colored by z-position showing (B) frontal and (C) ventral view of tracheoles, where anterior is violet and posterior is red. (D-E) Representative electron micrographs of brain neuropil showing regions that are (D) sparsely tracheated and (E) densely tracheated. Arrows indicate tracheoles that have been pseudocolored in pink. (F) Initial FlyWire segmentations associated with a manually traced tracheolar skeleton in CATMAID, showing erroneously merged neuronal and glial fragments (multiple colors). (G) Following manual proofreading, the same tracheole is displayed as an isolated, accurately segmented tracheole mesh. (H) Volumetric reconstruction in FlyWire showing the full, proofread, volumetric tracheolar network, with each tracheole represented as a unique color. Scale bars: 200 µm (A-C, H); 5 µm (D, E); 30 µm (F,G).

We next calculated the tracheole volume fraction within each brain synapse-rich super-region and neuropil subregion, as defined by (Ito et al., 2014) (Figure 2A, B). At the level of super-region, tracheole volume fraction differed substantially across anatomical divisions, ranging from 0.0281% in the lateral horn (LH) to 0.1405% in the central complex (CX) (Figure 2C, D). Across the 76 analyzed brain subregion neuropils, the median tracheole volume fraction was 0.0747%, with substantial anatomical variation across regions (Figure 2E, F).

**Figure 2.**
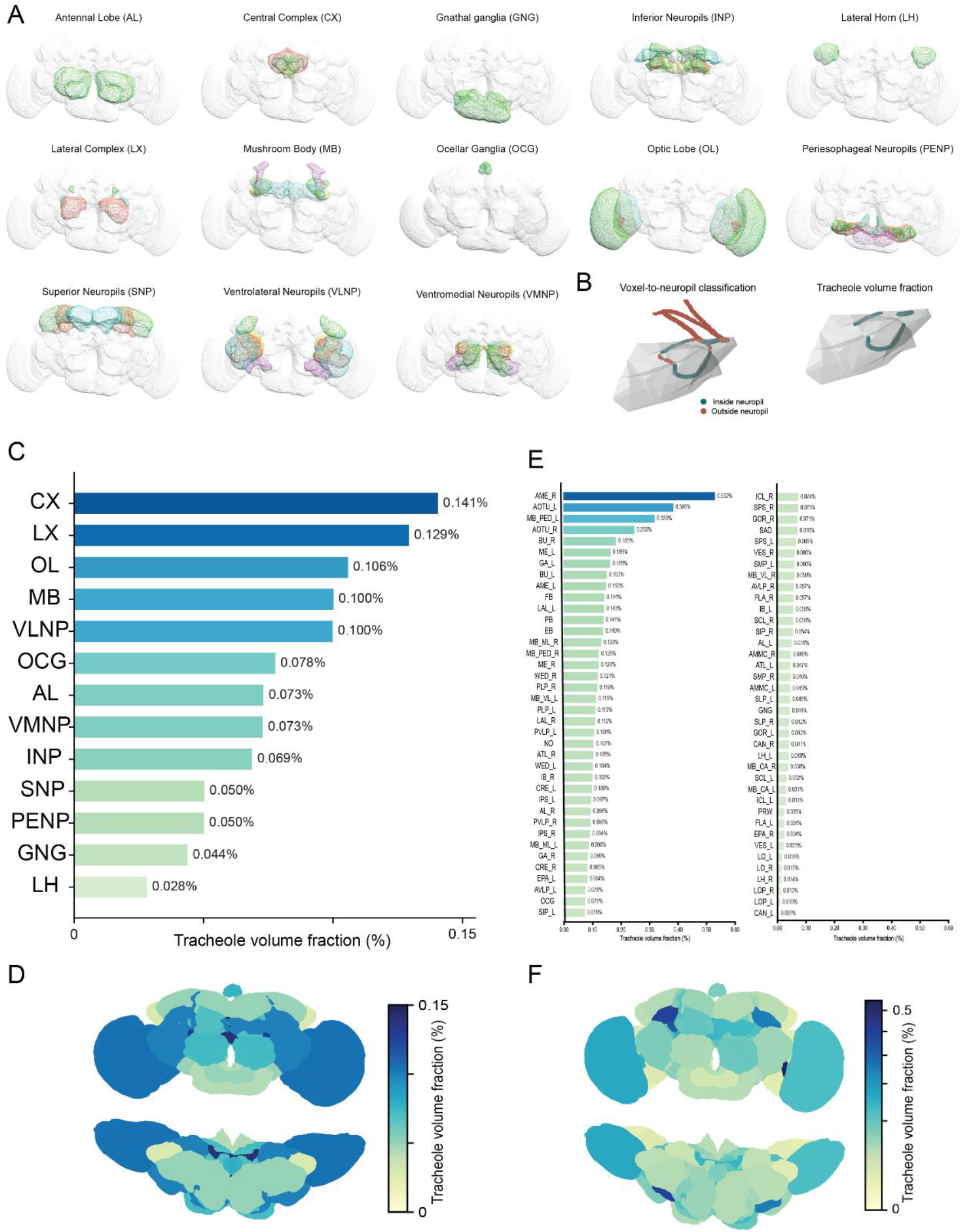
Tracheole volume fraction of synapse-rich neuropil volumes. (A) Schematic depicting the subvolumes that comprise each neuropil super-region (Ito et al., 2014). Super-regions (in bold) and their associated sub-regions are: **AL** = Antennal lobe (AL_L, AL_R); **CX** = Central complex (FB, EB, PB, NO); **GNG** = Gnathal ganglia; **INP** = Inferior neuropils (CRE_L, CRE_R, ICL_L, ICL_R, SCL_L, SCL_R, IB_L, IB_R, ATL_L, ATL_R); **LH** = Lateral horn (LH_L, LH_R); **LX** = Lateral complex (BU_L, BU_R, LAL_L, LAL_R, GA_L, GA_R); **MB** = Mushroom body (MB_CA_L, MB_CA_R, MB_PED_L, MB_PED_R, MB_VL_L, MB_VL_R, MB_ML_L, MB_ML_R); **OCG** = Ocellar ganglia; **OL** = Optic lobe (ME_L, ME_R, AME_L, AME_R, LO_L, LO_R, LOP_L, LOP_R); **PENP** = Periesophageal neuropils (SAD, AMMC_L, AMMC_R, FLA_L, FLA_R, CAN_L, CAN_R, PRW) ; **SNP** = Superior neuropils (SLP_L, SLP_R, SIP_L, SIP_R, SMP_L, SMP_R); **VLNP** = Ventrolateral neuropils (AOTU_L, AOTU_R, PVLP_L, PVLP_R, AVLP_L, AVLP_R, PLP_L, PLP_R, WED_L, WED_R); **VMNP** = Ventromedial neuropils (VES_L, VES_R, EPA_L, EPA_R, GOR_L, GOR_R, SPS_L, SPS_R, IPS_L, IPS_R). _L and _R denote left and right hemispheres. (B) Representation of a tracheole classified as either falling within (teal) or outside (orange) a defined neuropil volume mesh. Tracheole volume fraction was calculated by dividing the volume of tracheole mesh contained in the neuropil mesh volume by the neuropil volume, and multiplied by 100. (C) Tracheole volume fraction per synapse-rich neuropil super-region. Volume fraction differed across super-regions (Kruskal-Wallis H = 31.484, P < 0.001, ε² = 0.341). (D) Frontal and dorsal views of synapse-rich neuropil super-regions, color-coded by tracheole volume fraction. (E) Tracheole volume fraction per synapse-rich neuropil sub-region for all annotated neuropil subvolumes, ranked from highest to lowest. (F) Frontal and dorsal views of neuropil sub-regions colored by the tracheole volume fraction. Abbreviations: AME = Accessory medulla; AMMC = Antennal mechanosensory and motor center; AOTU = Anterior optic tubercle; ATL = Antler; AVLP = Anterior ventrolateral protocerebrum; BU = Bulb; CAN = Cantle; CRE = Crepine; EB = Ellipsoid body; EPA = Epaulette; FB = Fan-shaped body; FLA = Flange; GA = Gall; GNG = Gnathal ganglia; GOR = Gorget; IB = Inferior bridge; ICL = Inferior clamp; IPS = Inferior posterior slope; LAL = Lateral accessory lobe; LH = Lateral horn; LO = Lobula; LOP = Lobula plate; MB_CA = Mushroom body calyx; MB_ML = Mushroom body medial lobe; MB_PED = Mushroom body pedunculus; MB_VL = Mushroom body vertical lobe; ME = Medulla; NO = Noduli; PB = Protocerebral bridge; PLP = Posterior lateral protocerebrum; PRW = Prow; PVLP = Posterior ventrolateral protocerebrum; SAD = Saddle; SCL = Superior clamp; SIP = Superior intermediate protocerebrum; SLP = Superior lateral protocerebrum; SMP = Superior medial protocerebrum; SPS = Superior posterior slope; VES = Vest; WED = Wedge.

We then assessed left-right symmetry in tracheolar patterning by reflecting left-hemisphere tracheoles onto the right hemisphere and transforming the tracheoles within brain region-specific volumes to correct for differences in volume shape between the left and right brain (Figure 3A, B). Brain regions that spanned the midline were reflected across that midline. Region-specific rigid registration improved bilateral spatial correspondence, reducing the median spatial mismatch between mirrored tracheoles from 14.330 µm to 7.676 µm (Figure 3A, C). Even after registration, however, only about one third of tracheole centerline positions had a contralateral counterpart within 5 µm (Figure 3C). In contrast, neither total tracheole length nor tracheole volume fraction was consistently biased toward the left or right hemisphere across the bilateral neuropil pairs we examined (Figure 3C, D). Thus, each hemisphere receives a similar amount of tracheation, but the two sides are not precise mirror images of each other.

**Figure 3.**
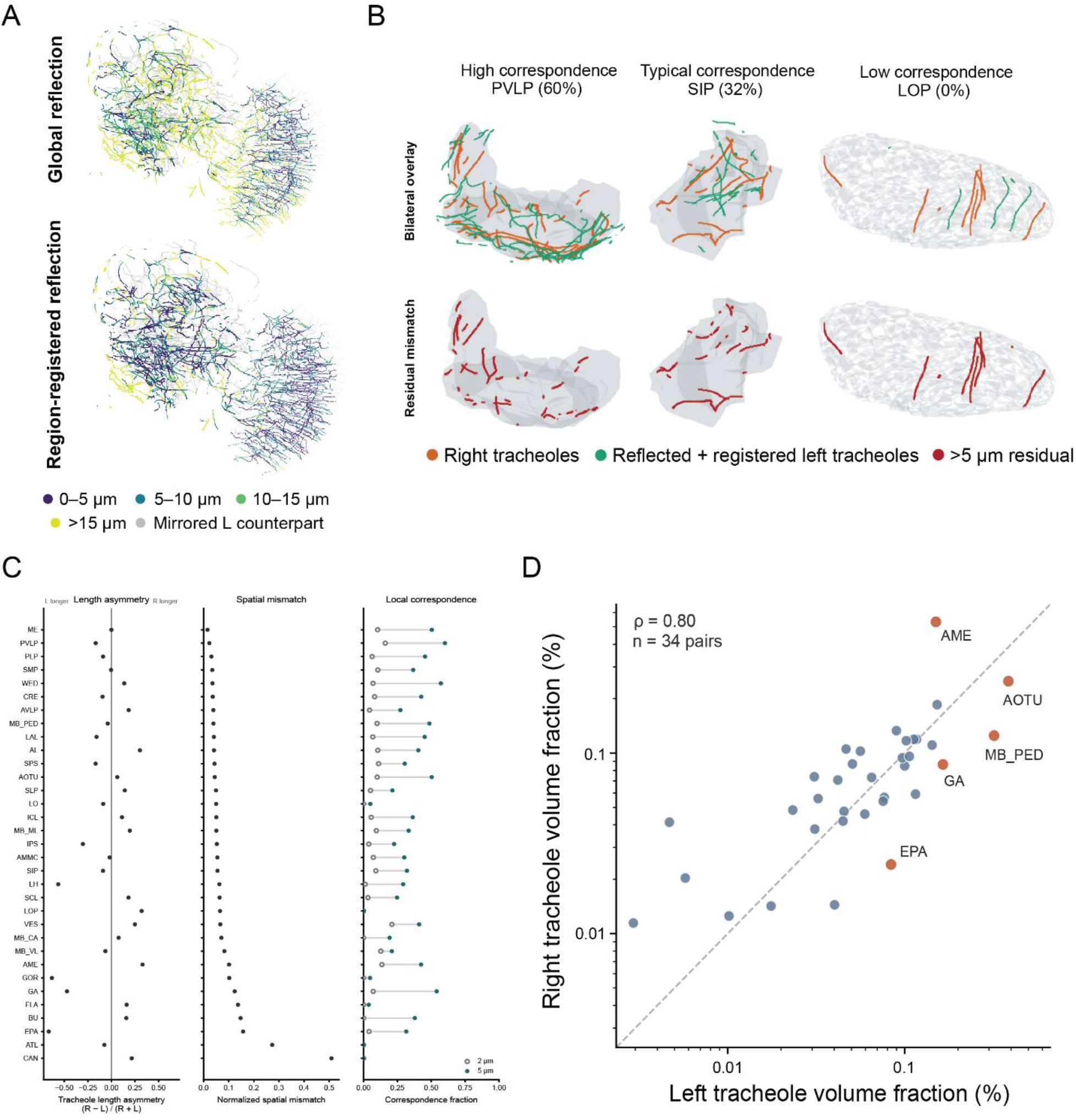
Bilateral symmetry of the tracheolar network. (A) Assessment of bilateral symmetry with right-hemisphere tracheoles shown with left-hemisphere tracheoles reflected across the midline by global reflection (top) or after registration of individual neuropil subregions (bottom). Tracheoles are colored by the distance from each node to the nearest node of its mirrored left counterpart. Across 33 neuropil pairs, region-specific registration reduced the median spatial mismatch from 14.330 µm to 7.676 µm (median reduction 5.980 µm, 95% CI 3.558-7.483 µm; Wilcoxon P < 0.001, paired rank-biserial = 0.986). After registration, the median normalized mismatch was 0.0528 (95% CI 0.0445-0.0654) and the median fraction of tracheole centerline positions with a contralateral counterpart within 5 µm was 0.3208 (95% CI 0.2723-0.4047). (B) Three representative neuropil volumes illustrating different levels of correspondence after reflecting and registering left-hemisphere tracheoles onto the right. Top row is the bilateral overlay of right tracheoles (orange) and reflected left-originating tracheoles (green). Bottom row is the same regions with mismatched tracheoles (> 5 µm) in red. Columns show a region of high correspondence between left and right, typical correspondence, and low correspondence. (C) Left, quantification of length asymmetry across bilateral neuropil pairs, calculated as (R-L)/(R+L). Positive values indicate greater length on the right, negative values indicate greater length on the left. Middle, normalized spatial mismatch between registered left tracheoles and existing right tracheole positions. Right, local correspondence between left and right showing the proportion of tracheole points with a matching point on the opposite side within 2 µm (open circles) or 5 µm (filled circles) after registration. (D) Tracheole volume fraction of right vs. left neuropil subregions (n = 34 paired subregions, Spearman ⍴= 0.80, P < 0.001). Volume fraction showed no systematic left-right bias (median right-minus-left difference +0.0022 percentage points, 95% CI -0.0136 to +0.0235; Wilcoxon P = 0.774, paired rank-biserial = 0.059); the signed asymmetry index (R-L)/(R+L) was centered near zero (median 0.0245, 95% CI -0.0824 to 0.1038, P = 0.566). Dashed line indicates perfect symmetry and orange points indicate subregions with the largest left-right disparities. Abbreviations as in Figure 2.

### Tracheoles enter the brain at discrete sites, branch extensively, and terminate within the brain parenchyma

The tracheolar network penetrated from outside the brain, through the neural lamella/glial sheath barrier, and into the brain at distinct locations we term infiltration points (Figure 4A-C). These infiltration points were predominantly located in the optic lobe (63/116) (Figure 4D). In contrast, no infiltration points were found in the central complex, lateral complex, or ocellar ganglion. Although the extra-brain origins of each tracheole cannot be definitively established due to the removal of air sacs during brain dissection, prior work indicates that the adult brain tracheoles arise from air sacs and cervical tracheae connected to thoracic spiracles (Chu et al., 2013; Wigglesworth, 1942). Surprisingly, there was one tracheole (1/117) that lacked an identifiable infiltration point, appearing to exist entirely within the brain, raising questions about its developmental origin and functional significance (Figure 4E, F; Supplemental Videos 2 and 3). Each tracheole branches extensively within the neuropil, and across all 117 tracheoles, the median distance from each traced point to the nearest point on a different tracheole was 8.55 µm (Figure 4G). Nearly all points (91.8%) along the tracheolar network were within 20 µm of a neighboring tracheole, and more than half (59.3%) were within 10 µm. Strahler analysis, a standard measure of branching complexity, revealed that most tracheoles have a Strahler number of 3 or 4 (93/117), while 23/117 tracheoles have a Strahler number of 1 or 2, and a single tracheole has a Strahler number of 5 (Figure 4H, I).

**Figure 4.**
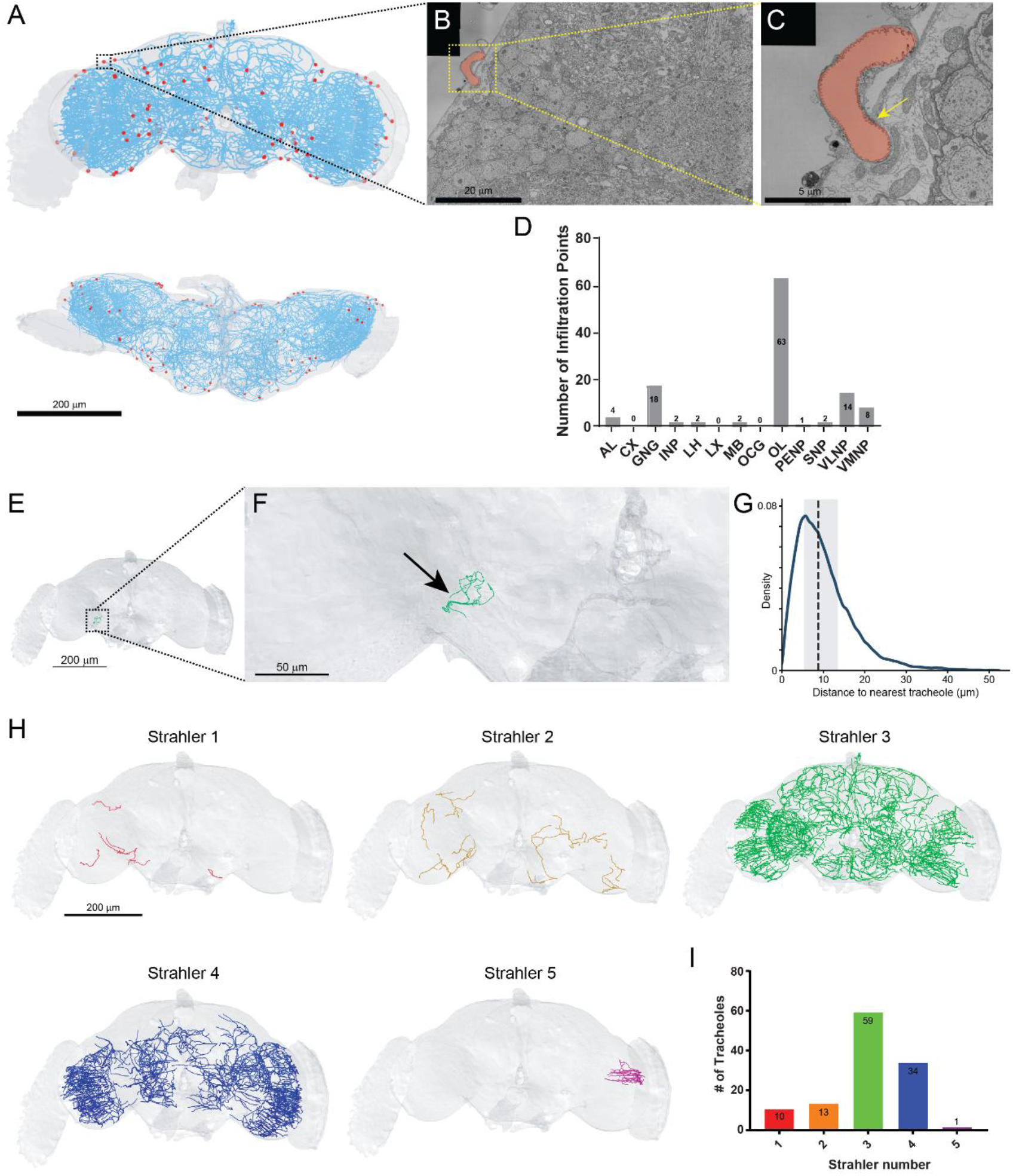
Infiltration and branching organization of individual tracheoles. (A) Frontal (top) and dorsal (bottom) 3D visualization of the skeletonized tracheolar network with individual infiltration points marked as red dots. (B) Representative electron micrographs showing tracheole infiltration into the brain parenchyma. (C) Higher magnification image of boxed area in B showing tracheole (pseudocolored pink) passing through the neural lamella/glial barrier (yellow arrow). (D) Total number of infiltration points per synapse-rich neuropil super-region in which they enter the brain. (E) 3D visualization of tracheole with no apparent infiltration point (green), overlaid on the FlyWire brain mesh (gray). The boxed region is enlarged in (F). (F) Higher magnification view of the tracheole (arrow), which lies entirely within the brain parenchyma and does not connect to the brain surface. See also Supplemental Videos 2 and 3. (G) Kernel density estimate of the three-dimensional distance from each CATMAID tracheole node to the nearest node belonging to a different tracheole. Median distance (dashed line) = 8.55 µm; shading indicates the interquartile range (5.16-13.14 µm). (H) 3D visualization of all tracheoles classified according to their branching complexity and Strahler number. (I) Number of tracheoles at each Strahler number. Scale bars: 200 μm (A, E, H); 50 μm (F); 20 μm (B); 5 μm (C). Abbreviations as in Figure 2.

We find that the brain’s tracheolar network is not a continuous reticular system, as each tracheole terminates within the brain parenchyma. In total, the 117 tracheoles culminate in 1,376 discrete intraparenchymal terminal ends that typically end via membranous compaction (Figure 5A, B; Supplemental Video 4). However, unlike the electron-lucent tracheolar lumen (presumably gas-filled, *in vivo*), the lumen of tracheolar ends were frequently somewhat electron-dense and contained vesicle-like structures (Figure 5C, D), perhaps indicating the presence of fluid within tracheolar endings.

**Figure 5.**
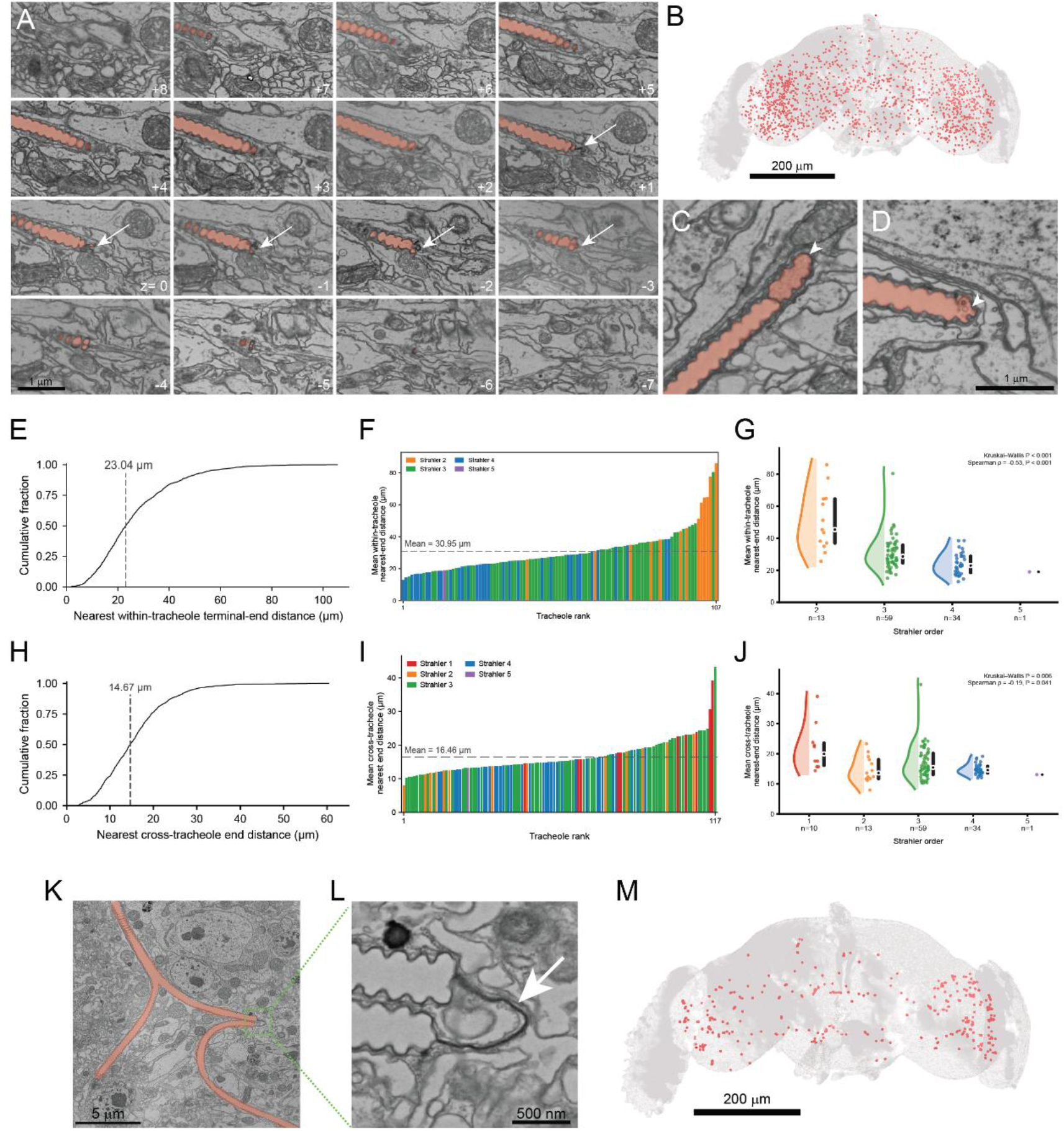
Intra-parenchymal tracheole endings and membranous tethers. (A) Serial electron micrographs (16 consecutive z-sections) showing a single tracheole ending within the brain parenchyma. Arrows indicate regions of membranous compaction at the distal end of the tracheole lumen (pseudocolored pink). (B) Three-dimensional visualization of 1,376 tracheole terminal ends (red dots) distributed throughout the brain. (C-D) Examples of tracheole endings (arrowheads) showing: (C) a distal apparently fluid-filled compartment and (D) membranous structures within the tracheole lumen. (E) Cumulative distribution of nearest terminal-end distances within the same tracheole. Pooled terminal-end median = 23.04 µm. Tracheoles containing only one terminal end were excluded. (F) Mean within-tracheole nearest-end distance for each tracheole, ranked in ascending order and colored by Strahler number. (G) Distribution of tracheole-level mean within-tracheole nearest-end distance by Strahler number. The single Strahler-5 tracheole is shown descriptively but was excluded from inferential analysis; across Strahler numbers 2-4, Kruskal-Wallis H(2) = 30.974, P < 0.001; Spearman ρ = -0.527, 95% CI -0.657 to -0.371, P < 0.001; pairwise comparisons for Strahler numbers 2 vs 3, 2 vs 4 and 3 vs 4 all q < 0.001 after FDR correction. (H) Cumulative distribution of nearest terminal-end distances between different tracheoles. Pooled terminal-end median = 14.67 µm. (I) Mean cross-tracheole nearest-end distance for each tracheole, ranked in ascending order and colored by Strahler number. (J) Distribution of tracheole-level mean cross-tracheole nearest-end distance by Strahler number. The single Strahler-5 tracheole is shown descriptively but was excluded from inferential analysis; across Strahler numbers 1-4, Kruskal-Wallis H(3) = 12.459, P = 0.006; Spearman ρ = -0.190, 95% CI -0.356 to -0.012, P = 0.041. (K) Electron micrograph showing a tracheole link connector (TLC), a membranous tether connecting two adjacent tracheole lumens (pseudocolored pink), characterized by a compacted, non-luminal membrane at the junction (dashed box). (L) Magnification of boxed region showing a TLC (arrow). (M) Three-dimensional visualization of 386 TLCs distributed throughout the brain. Scale bars 1 μm (A, C, D); 200 μm (B, M); 5 μm (K); 500 nm (L).

To characterize how these endings are spatially organized, we first asked how close together endings are on the same tracheole. The median within-tracheole nearest end distance was 23.04 µm, though some end pairs were separated by more than 100 µm (Figure 5E). We then asked whether this spacing depends on how extensively a tracheole branches. Grouping tracheoles by Strahler number, we found that the more a tracheole branches, the closer together its endings are positioned (Figure 5F, G). We next asked whether endings on different tracheoles show a similar relationship, by measuring the distance from each ending to the nearest ending on a different tracheole (Figure 5H). We found that more complex tracheoles also had closer cross-tracheole endings (Figure 5I, J). Together, these data suggest that as tracheoles branch more extensively, their endings are found closer together, though the extent to which neighboring tracheoles influence the spacing of individual tracheole endings remains an open question.

While characterizing the terminal tracheole endings, we identified numerous novel membranous tethers that we termed tracheole link connectors (TLC) that appear to be local areas of tracheolar collapse or compaction, where gaseous luminal space is absent. The TLCs seem not to be *bona fide* tracheolar endings, as tracheole lumen continues on both sides of the TLC (Figure 5K, L; Supplemental Video 5). We identified 386 unique TLCs distributed throughout the brain (Figure 5M). Additional experiments are necessary to determine whether these membranous tethers are physiologically relevant to respiration or whether they are artifacts of fixation/processing.

### Synapse proximity to tracheoles varies by brain region

Synaptic transmission is energetically demanding, and presynaptic terminals meet this demand primarily through oxidative phosphorylation in mitochondria (Justs et al., 2022). To examine how tracheoles are positioned relative to these sites of high oxygen demand, we measured the distance from each synapse to the nearest tracheole using recently published synapse annotations (Yu et al., 2025). Tracheoles were frequently observed within synapse-rich neuropil (Figure 6A); across 78.3 million presynaptic contacts in 76 brain neuropils, the median distance to the nearest tracheole skeleton was 7.87 µm, and nearly every synapse (97.3%) was located within 20 µm of a tracheole. However, median synapse-to-tracheole proximity differed across the 13 super-regions, with regional median distances ranging from 6.47 to 12.36 µm (Figure 6B). We observed a similar regional pattern for each neurotransmitter, with serotonergic and glutamatergic synapses varying the most across brain regions (Figure 6C, D).

**Figure 6.**
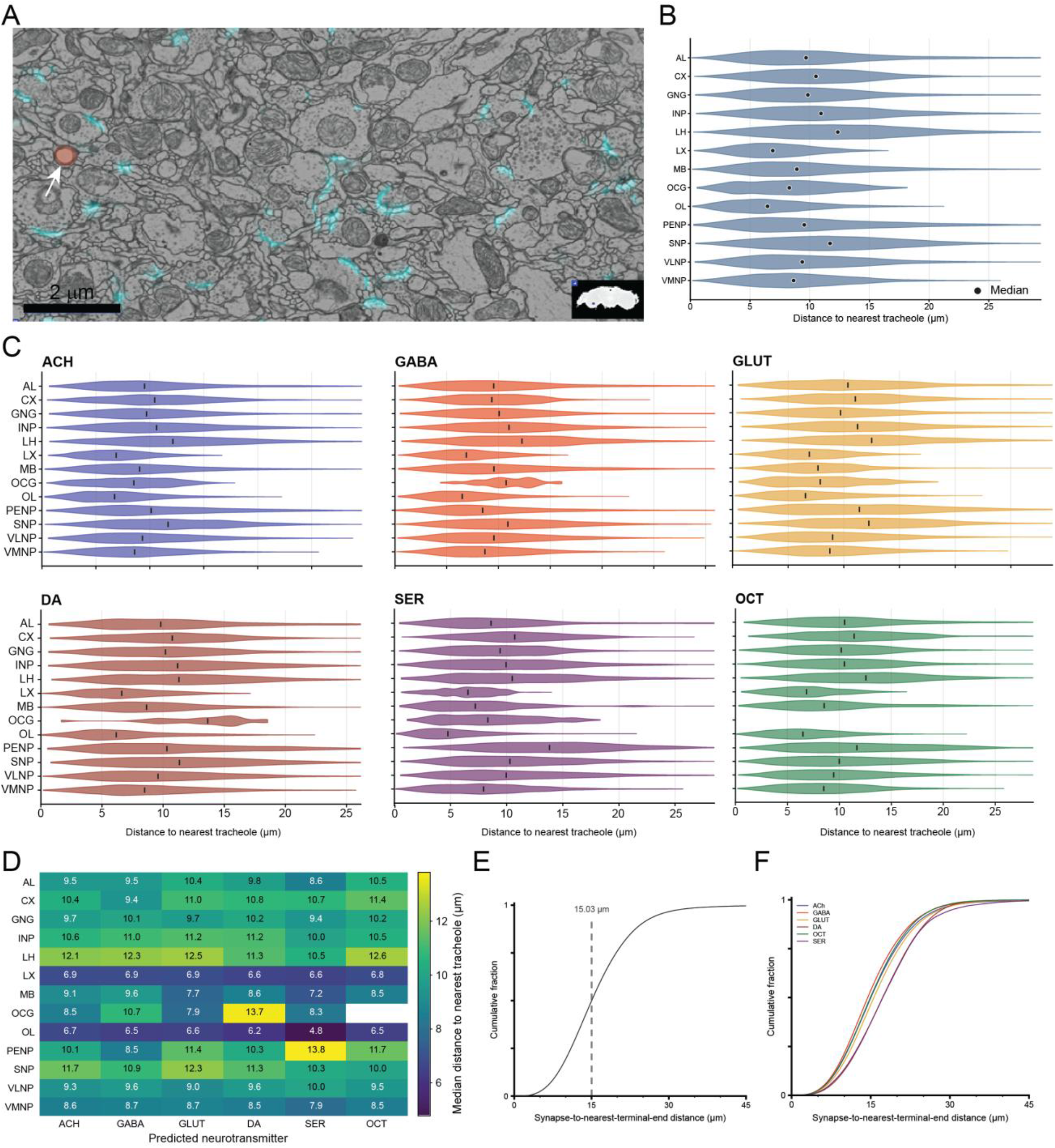
Proximity of presynaptic contacts to brain tracheoles. (A) Electron micrograph showing a tracheole in cross section (pseudocolored in pink, white arrow) surrounded by predicted synapses (cyan). (B) Distributions of the distance from presynaptic contacts to the nearest sampled tracheole centerline point across synapse-rich neuropil super-regions. Distances differed across the 13 super-regions (Kruskal-Wallis H = 13,876.5, P < 0.001, ε² = 0.107). (C) The same analysis stratified by predicted neurotransmitter class: acetylcholine (ACH), GABA, glutamate (GLUT), dopamine (DA), serotonin (SER), and octopamine (OCT). Vertical ticks indicate medians. Regional dependence was retained within every neurotransmitter class (ε² = 0.102-0.174; strongest for SER, ε² = 0.174, followed by GLUT, ε² = 0.138). (D) Median synapse-to-tracheole centerline distance for each super-region and predicted neurotransmitter combination. The OCG/OCT cell is empty because no octopaminergic contacts in that region met the neurotransmitter-confidence threshold. (E) Cumulative distribution of the distance from presynaptic contacts to the nearest tracheole terminal end. Median = 15.03 µm. (F) Terminal-end distance distributions stratified by predicted neurotransmitter class. Distance distributions differed across neurotransmitter classes (χ²(250) = 336,836, P < 0.001, Cramér’s V = 0.031). Scale bar: 2 µm (A). Abbreviations as in Figure 2.

Tracheolar terminal ends have been shown to be recruited by hypoxic tissue (Jarecki et al., 1999), so they might be preferentially positioned near synapses if synapses drive local oxygen demand. We therefore measured the distance from each presynaptic contact to the nearest tracheole terminal end (Figure 6E). The median distance was 15.03 µm, approximately twice the median distance to the nearest point on tracheole skeletons (7.87 µm). Stratifying by neurotransmitter, GABAergic contacts were closest to terminal ends, whereas serotonergic and dopaminergic contacts were farthest (Figure 6F).

### Synapse proximity to tracheoles reveals neurotransmitter- and region-specific heterogeneity

The proximity of synapses to tracheoles may be determined by a biological relationship, or it could be that synapses found in regions of high tracheolar fraction appear closer to tracheoles simply because there are more tracheoles present. To distinguish between these possibilities, we compared the median distance from each neurotransmitter-specific synapse population to a null distribution built by resampling an equal number of synapses from the same region, independent of neurotransmitter identity (see Methods). Relative to this regional expectation, acetylcholinergic and glutamatergic synapses were slightly farther from tracheoles, GABAergic, dopaminergic, and serotonergic synapses were closer, and octopaminergic synapses did not show a whole-brain bias (Figure 7A). These effects varied considerably across brain regions, with 69 of 77 region/neurotransmitter combinations reaching significance after correction for multiple comparisons (Figure 7A, B).

**Figure 7.**
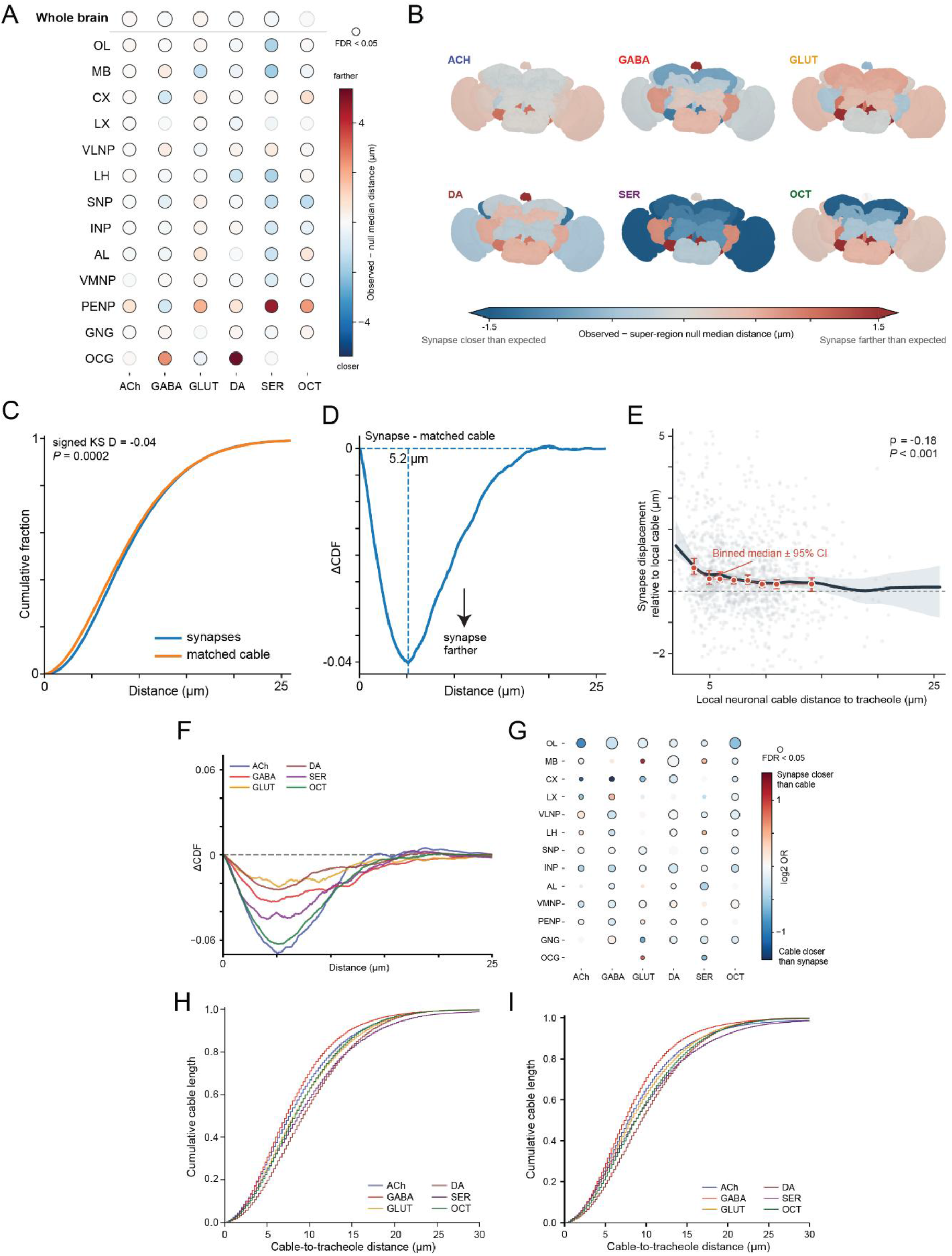
Synapse proximity to tracheoles and displacement of synapses from matched neuronal cable. (A) Neurotransmitter-specific synapse-to-tracheole proximity relative to anatomical expectation. The top row shows the anatomy-matched whole-brain effect, for which each neurotransmitter’s observed contact count was preserved separately within each super-region before null samples were pooled across the brain. Remaining rows show region-specific effects relative to null samples drawn from all presynaptic contacts within the same super-region. Values represent the observed median minus the null median. Whole-brain effects were ACh +0.104 µm, GLUT +0.270 µm, GABA -0.045 µm, DA -0.100 µm, SER -0.194 µm and OCT +0.007 µm; all except OCT remained significant after FDR correction (q < 0.001). Bold circles indicate FDR-adjusted *q* < 0.05. (B) Region-specific effects from (A) rendered anatomically for each neurotransmitter class. (C) Matched cumulative distribution of distance to the nearest tracheole surface for presynaptic sites (blue) and matched non-synaptic cable controls from the same neurons (orange). Median presynaptic distance was 8.42 µm versus 8.12 µm for matched cable; synapses were farther from tracheoles than matched cable overall (signed KS D = −0.04; conditional logistic regression OR = 0.884 per 5 µm closer to the tracheole surface, P < 0.001), and at the neuron level the median synaptic displacement was +0.36 µm (Wilcoxon P < 0.0001). (D) Difference between the two curves in (C). Negative values indicate synapses are farther from tracheoles than matched cable at the same distance, with the largest deviation occurring at 5.2 μm from the tracheole. (E) Relationship between local matched-cable distance to the tracheole surface and site-level synaptic displacement relative to matched cable (Spearman ρ = -0.18, P < 0.001). Gray points represent individual presynaptic sites; red points and error bars show binned medians and 95% confidence intervals; the black line and shaded band show a LOWESS fit with bootstrap confidence intervals. (F) ΔECDF as in (D), computed separately for each predicted neurotransmitter. (G) Odds ratio of a point being synaptic, relative to matched cable, for each super-region and predicted neurotransmitter, expressed as log2 OR; 60 of 74 region/neurotransmitter combinations reached significance. Bold circles indicate FDR-adjusted *q* < 0.05. (H) Cable-length-weighted cumulative distributions of presynaptic-arbor cable distance to the nearest tracheole mesh surface, stratified by predicted neurotransmitter class. Formal comparisons used one neuron as the statistical unit (ACh, n = 242; GABA, n = 248; GLUT, n = 248; DA, n = 249; SER, n = 240; OCT, n = 42). Cable-to-tracheole proximity differed across neurotransmitter classes (Kruskal-Wallis H(5) = 236.445, P < 0.001, ε² = 0.183). (I) Cumulative distribution of the distance from neuronal cable to the nearest tracheole mesh surface, weighted by cable length and stratified by predicted neurotransmitter, in which the entire neuronal skeleton, rather than the presynaptic-arbor subtree, was used to measure cable-to-tracheole distance. Abbreviations as in Figure 2.

### Synapses sit farther from tracheoles than matched neuronal cable

The previous analysis accounts for differences in how densely tracheated each brain region is, but not for where individual neurons travel within a region. A synapse could lie close to a tracheole because synapses themselves are preferentially positioned near the tracheolar network, or simply because the neuron forming it happens to run near a tracheole. To separate these two possibilities, we measured how far each synapse was from the nearest tracheole and compared it with points on the same neuron without synapses. If synapses are specifically positioned near tracheoles, they should be closer than the rest of the neuron, and if not, the two distances should be similar. We found that synapses were not closer to tracheoles than the rest of their neuron. Instead, synapses sat slightly farther away, with a median shift of 0.36 µm away from the tracheole (Figure 7C). The difference between synapses and cable was greatest about 5 µm from a tracheole and was not observed beyond approximately 15 µm (Figure 7D), suggesting that this effect is restricted to the space immediately surrounding tracheoles. Consistent with this, the shift was strongest where a neuron passed within a few micrometers of a tracheole (Figure 7E). All six neurotransmitter classes showed the same overall direction, although by different amounts (Figure 7F). The size of the shift also varied across brain regions, with most region and neurotransmitter combinations showing small but statistically significant effects (Figure 7G).

Because synapses are located on neuronal branches, we next asked whether the neurotransmitter-specific differences we observed could simply reflect where these neurons are positioned relative to tracheoles. To test this, we measured the distance from each neuron’s presynaptic branches to the nearest tracheole, independent of where its synapses are located.

We found that the presynaptic arbors of different neurotransmitter classes occupy different distances from the tracheolar network (Figure 7H), and this held true when we analyzed entire neuronal skeletons, as well (Figure 7I). Together, these analyses suggest that neurotransmitter classes differ in where their branches sit relative to tracheoles, but differ only modestly in where synapses form along those branches. Indeed, for every neurotransmitter class, synapses were positioned slightly farther from tracheoles than the rest of the neuron’s own cable.

## Discussion

This study presents a comprehensive neuroanatomical characterization of the tracheolar network within a female adult *Drosophila melanogaster* brain. While the tracheal systems of embryonic, larval, and adult thoracic and abdominal regions have been extensively studied (Bossen et al., 2023; Ehrhardt et al., 2022; Hayashi & Kondo, 2018), the intricate architecture of the adult brain tracheolar network remains less well understood.

The brain’s tracheolar system originates from the cervical trachea, an extension of the most anterior thoracic spiracle from within the first segmental tracheal network (Hartenstein et al., 2008; Pereanu et al., 2007). We find that this tracheolar network infiltrates the brain through distinct entry points (Figure 4). The distribution of tracheolar entry points is notably non-uniform, with a substantial enrichment within the optic lobes. This raises questions concerning the factors that govern tracheolar patterning. Notably, the optic lobes represent the largest neuropil structures in the adult fly brain, and 54% of the brain’s ∼140,000 neurons are intrinsic to the optic lobes (Schlegel et al., 2024). These neurons are devoted to visual processing and therefore likely impose substantial metabolic demands (Laughlin et al., 1998; Malin & Desplan, 2021). Moreover, anatomical proximity may also contribute to the observed pattern. The optic lobes occupy the lateral and posterior surfaces of the brain, immediately adjacent to four of the six major head air sacs through which tracheoles enter the brain (Chu et al., 2013; Whitten, 1957). The high density of tracheolar infiltration into the optic lobes may thus reflect their location as the most accessible interface between the brain parenchyma and the surrounding air sac/tracheal network. Future functional studies comparing neuronal activity and oxygen consumption with regional tracheole volume fraction will be necessary to determine the relative contributions of metabolic demand and/or developmental anatomy to tracheolar patterning.

From these infiltration points, tracheoles branch extensively within the brain parenchyma, terminating in discrete terminal ends, without forming an interconnected vascular-like network. This structural organization suggests a reliance on diffusion-based gas exchange (Lehmann & Schützner, 2010; Pickard, 1974), in contrast to the circulatory systems of vertebrates (Willie et al., 2014; Wu et al., 2022). Because diffusion is effective only over short distances, and because each tracheole ends on its own rather than connecting with its neighbors, where tracheoles terminate may be particularly important for meeting the local metabolic needs of each brain region (Jarecki et al., 1999). Indeed, the terminal endings of tracheoles sometimes appeared to contain fluid (Figure 5C, D), which is notable given that fluid within distal tracheoles has been proposed to regulate activity-dependent diffusion of gas into tracheated tissue (Wigglesworth, 1930; Wigglesworth & Lee, 1982). One likely contributor to brain tracheolar patterning is the Breathless/Branchless (FGFR/FGF) signaling pathway, which plays a crucial role in early tracheal patterning throughout the fly body (Klämbt et al., 1992; Sutherland et al., 1996). Tracheole ingrowth into the adult visual system depends on this pathway during metamorphosis (Chu et al., 2013), and whether comparable mechanisms contribute to the end patterning of this network in adults remains to be determined. Notably, we identified a tracheole that is completely embedded within the parenchyma of this FAFB dataset, with no apparent infiltration point (Figure 4E, F; Supplemental Videos 2 and 3). The developmental origin and functional relevance of a tracheole disconnected from the spiracular-originating tracheal network remains unclear. Its existence could simply be related to a fixation or tissue processing artifact, or it could be a branch that has become separated from the tracheolar network.

The six air sacs surrounding the brain are hypothesized to facilitate the movement of air through the tracheolar network (Chu et al., 2013; Whitten, 1957). The presence of tracheole regions devoid of lumen, which we term tracheole link connectors (TLC), raises questions about how gas would flow through these tracheolar sections. While tracheal compressions have been documented during synchrotron monitoring of tracheal tubes in the head of beetles, even during maximal compression, there was still tracheal lumen present (Westneat et al., 2003). It has been hypothesized that tracheal migration and growth are directed by long filaments that extend forth from epidermal cells associated with trachea (Wigglesworth, 1959). Whether the observed TLCs are related to filamentous protrusions associated with tracheole growth, or whether they are fixation artifacts introduced during sample processing, will require additional investigation. An additional possibility is that TLCs represent specialized sites involved in regulating local gas movement within the tracheolar network. In many insects, gas exchange is dynamically regulated through spiracular activity and continuous, cyclic, or discontinuous gas exchange cycles, which generate oxygen and carbon dioxide flux throughout the tracheal network (Gefen & Matthews, 2021; Matthews, 2018). The localized loss of lumen observed at TLCs raises the possibility that these structures influence local gas movement or compartmentalize regions of the tracheolar network. Future studies examining tracheolar morphology during respiration will be necessary to determine the physiological relevance, if any, of TLCs.

Neurons depend on a continuous supply of oxygen to meet the metabolic demands of synaptic transmission and neuronal physiology. Their energy needs are met largely by ATP produced through oxidative phosphorylation in mitochondria, and in *Drosophila*, presynaptic terminals rely primarily on this mitochondrial pathway (Justs et al., 2022). To characterize the spatial relationship between tracheoles and synapses, we compared each synapse with matched non-synaptic cable from the same neuron. Synapses were displaced away from the immediate vicinity of tracheoles relative to their own neuron’s surrounding cable (Figure 7C, D), and this displacement was strongest where a neuron’s processes ran closest to a tracheole (Figure 7E). One possible explanation is that oxygen use is set by the location of mitochondria rather than synapses. If mitochondria along axonal segments adjacent to tracheoles supply sufficient ATP to nearby synapses, there may be little pressure for the synapses themselves to sit close to a tracheole. As mitochondrial segmentations become more widely available (Sager et al., 2026), quantifying mitochondria along axons and at synapses will help determine the relationship between tracheolar patterning, axonal mitochondrial distribution, and synapse placement.

A second possible contributor to the relative depletion of synapses near tracheoles is the presence of surrounding glial cells. Glia are known to constrain tracheole growth during development (Pereanu et al., 2007), and in our tracheolar reconstruction, we frequently found glial cell processes directly adjacent to tracheoles, which had to be manually separated from the tracheolar meshes during proofreading (Figure 1F, G). Relatedly, *Drosophila* glia envelop nearly all neuronal compartments (Kremer et al., 2017), astrocytic membranes associate closely with tracheoles and with synapses (Freeman, 2015), and barrier-associated glia extend processes into the neuropil interior (Contreras et al., 2024). Whether these or other glial cell subtypes specifically contact tracheoles remains unknown. Unlike neurons, glia have not been systematically reconstructed in most whole-brain EM datasets, including FAFB, because their thin sheet-like processes are difficult to segment. Resolving their contribution to local tracheolar patterning and synaptic exclusion will therefore require more reliable glial reconstructions, in addition to functional glial-tracheole studies.

We find the brain’s tracheolar system to be largely symmetric in how much tracheation each hemisphere receives, but the individual branching within brain region compartments does not mirror the opposite side (Figure 3). Future studies extending this work to additional brain volumes will be required to quantify variability across animals. Additionally, sex-specific differences in tracheal morphology have been documented in the *Drosophila* gut, where females show greater tracheal length and branching than males (Blackie et al., 2024). Reconstructing additional brains of both sexes will be necessary to better understand if there are sex-dependent differences or general rules to brain tracheolar patterning.

## Conclusion

We have mapped the complete tracheolar network of an adult *Drosophila* brain. This network is made up of independently terminating tracheoles that enter the brain at discrete infiltration points and end within the parenchyma, without forming an interconnected, vascular-like network. Tracheoles are heterogeneously distributed across neuropils, and are largely symmetric between the two hemispheres in overall tracheation, though not in their precise branching patterns. We also find that synapses are not positioned closer to tracheoles than the neuronal branches they arise from. Together, this work provides a foundation for future studies examining the dynamic interplay between neuronal activity and metabolic demand in the *Drosophila* brain, and serves as a reference for comparing tracheolar organization across other volumetric electron microscopy brain datasets.

## Supporting Information

Supplemental video files can be found at: https://drive.google.com/drive/u/0/folders/102bdayfPmrHNOBW-JVWTGurmsJ7ETrLM

**Supplemental Video 1. Depth-mapped tracheoles.** Rotating frontal view of all tracheoles rainbow colored by z-position, where anterior is violet and posterior is red.

**Supplemental Video 2. Tracheole without an infiltration point in 3D.** Rotating, oblique view of FlyWire mesh (gray), showing the tracheole with no apparent infiltration point (green) and its position within the brain parenchyma.

**Supplemental Video 3. Tracheole without an infiltration point in raw EM images.** Animation stepping through raw electron microscopy images in the z direction. The tracheole (pseudocolored green) is contained within the brain, and remains separated from the neural lamella throughout, with no discernible infiltration point.

**Supplemental Video 4. Tracheole ending.** Animation stepping through raw electron microscopy images in the z direction, showing the terminal end of a tracheole within the neuropil (pseudocolored pink).

**Supplemental Video 5. Tracheole link connector (TLC).** Animation stepping through raw electron microscopy images in the z direction, showing a TLC tether without a lumen that connects two parts of a lumen-containing tracheole (pseudocolored light blue).

## Acknowledgements

We thank Tom Kazimiers (Kazmos GmbH) for expert technical advice with CATMAID. We also thank the Princeton FlyWire team and members of the Murthy and Seung labs, as well as members of the Allen Institute for Brain Science, for development and maintenance of FlyWire (supported by BRAIN Initiative grants MH117815 and NS126935 to Murthy and Seung). We also acknowledge members of the Princeton FlyWire team and the FlyWire consortium for neuron proofreading and annotation.

## Author contributions

Conceptualization (KJC); Data Curation (KS, HC, TN, SH, JS, AM, SL, BW, RSK, MB, LZ, FR-C, EA, YC, CAL, KB, KL, CM); Formal Analysis (KJC, DDB, PS, AMD, HC, TN, KS, JS, AM, SL, MB); Funding acquisition (KJC, AMD, PS); Investigation (KJC, DDB, PS, AMD, HC, TN); Methodology (KJC, DDB, PS, AMD, KS, HC, TN); Project Administration (KJC, KS); Resources (DDB, AMD, PS); Software (DDB, AMD, PS); Supervision (KJC); Validation (KJC, KS); Visualization (KJC, PS, HC, TN, KS, JS); Writing – original draft (KJC, HC, TN); Writing – review and editing (all authors).

## Data availability

Data and source code will be stored on GitHub, and will be made publicly available upon acceptance. Editors and reviewers can obtain them on request.

